# Investigating densities of Symbiodiniaceae in two species of Antipatharians (black corals) from Madagascar

**DOI:** 10.1101/2021.01.22.427691

**Authors:** Erika Gress, Igor Eeckhaut, Mathilde Godefroid, Philippe Dubois, Jonathan Richir, Lucas Terrana

## Abstract

Here, we report the first methodological approach to investigate the presence and estimate the density of Symbiodiniaceae cells in corals of the order Antipatharia subclass Hexacorallia, known as black corals. Antipatharians are understudied ecosystem engineers of shallow (<30 m depth), mesophotic (30-150 m) and deep-sea (>200 m) reefs. They provide habitat to a vast number of marine fauna, enhancing and supporting coral reefs biodiversity globally. Nonetheless, little biological and ecological information exists on antipatharians, including the extent at which global change disturbances are threatening these corals. The assumption that they were exempted from threats related to climate change was challenged by findings of high density of dinoflagellates within three antipatharian colonies. Further methodical studies were necessary to investigate the regularity of these findings. An integrated design combining microscopy and molecular techniques was used to investigate the presence and estimate density of Symbiodiniaceae cells within two antipatharians species -*Cupressopathes abies* and *Stichopathes maldivensis* -from shallow and mesophotic reefs of SW Madagascar. Symbiodiniaceae-like cells were found within the two species from both shallow and mesophotic reefs, although the overall cell density was very low (0-4 cell mm^-3^). These findings suggest that high abundance of Symbiodiniaceae is not characteristic of antipatharians, which has relevant implications considering disruptions associated to climate change affecting other corals. However, the high densities of dinoflagellates found in antipatharian colonies exposed to higher light irradiance in other studies should be further examined.

## 1. Introduction

Coral reefs are the most biodiverse marine ecosystems and are among the most biologically rich and productive ecosystems on Earth (Rogers 1979). However, most of this information is based on our knowledge of shallow reefs (<30 m depth). Light-dependant reefs can thrive in deeper waters though. Mesophotic coral ecosystems (MCEs) are light-dependant coral reefs found typically from 30 to 150 m in tropical and subtropical regions (Hinderstein et al. 2010). Hereinafter MCEs also referred as ‘mesophotic reefs’ for the ease of the reader and to be analogous with the term ‘shallow reefs’. Mesophotic reefs have been estimated to represent about 80% of potential coral reef habitat worldwide – still, we know little about them compared to shallow reefs (Pyle and Copus 2019). Scleractinians (hard corals) are present in MCEs, but to a much lesser extent than in shallow reefs (Baker et al. 2016; Gress et al. 2018; Pyle and Copus 2019). Antipatharians, octocorals, sponges and macroalgae provide most of the available habitat on mesophotic reefs (Baker et al. 2016; Gress et al. 2018; Pyle and Copus 2019).

The order Antipatharia (subclass Hexacorallia) comprises seven families and around 270 species (Brugler et al. 2013; Opresko 2019). These corals occur in most oceans at depths ranging from 2 to 8,900 m, and are known to favour strong currents and low-light environments (Grigg 1965; Wagner et al. 2012). Antipatharians do not produce a calcium carbonate skeleton; instead, the thorny axial skeleton is composed of a proteinaceous complex called antipathin (Goldberg 1978; Goldberg et al. 1994). They tend to increase in diversity and abundance with depth, reaching a peak at mesophotic depths (30 -150 m) (Bo et al. 2019). Nonetheless, dense aggregations have also been observed in shallow (<30 m depth) clear water environments (Tazioli et al. 2007; Suarez et al. 2015; Gress pers. obs.). Antipatharians are crucial habitat-providing corals with which an array of marine fauna associate (Parrish et al. 2002; Boland and Parrish 2005; Tazioli et al. 2007; Wagner et al. 2012; Bruckner 2016; Terrana et al. 2019). For instance, in the Philippines the average density of invertebrate macrofauna associated with antipatharians ranged from 82 to 8,313 individuals/m^2^ (Suarez et al. 2015). Therefore, antipatharians can be considered ecosystem engineers supporting and enhancing marine biodiversity at shallow, mesophotic and deep-sea depths. However, limited studies have evaluated their current condition under increasing threats, such as those associated with climate change.

Disturbance to the relationship between dinoflagellates and scleractinians – on shallow and mesophotic reefs – due to climate change is an increasing threat (Baker et al. 2008; Hughes et al. 2018; Bongaerts and Smith 2019; Fisher et al. 2019). Nevertheless, our current understanding of the physiological mechanisms underlying the endosymbiotic association between dinoflagellates and their cnidarian host has been constantly revised and is still not fully understood. To support further discussion of the present study findings, in combination with findings from previous studies, a summary of the coral-algae known mechanisms is appropriate. Four of the seven genera of Symbiodiniaceae are found in symbioses with scleractinian corals (*Symbiodinium, Breviolum, Cladocopium* and *Durusdinium*) (LaJeunesse et al. 2018; Meistertzheim et al. 2019). The translocation of photosynthetically fixed carbon from the symbiont to the host is considered the best-known aspect of the coral-algae symbiosis. However, the amount of photosynthetic carbon translocated to the host and the identity of the compounds are not fully known (Davy et al. 2012). The resulting lack or low number of dinoflagellates and their photosynthetic pigments leads to ‘bleaching’, which occurs due to oxidative stress. During stressful conditions, three potential mechanisms have been suggested for the regulation of symbiont numbers in alga-invertebrate symbioses: (i) expulsion of excess symbionts; (ii) degradation of symbionts by host cells; and (iii) inhibition of symbiont cell growth and division controlled by the pH of the host cell (Davy et al. 2012; Barott et al. 2015).

In marine environments, oxidative stress results from the production and accumulation of reactive oxygen species (ROS) – as a response to heat, ultraviolet radiation and pollution – and can damage lipids, proteins and DNA (Lesser 2006; Roth 2014). Over prolonged periods of water temperature alterations, dinoflagellates can become a substantial source of ROS that is transferred to and accumulated in the host (Lesser 2006; Weis 2008; Roth 2014). When corals reduce the number of their symbionts, the main source of ROS production is removed, although the coral host itself may also produce ROS as a result of light and temperature (Roth 2014). High concentrations of ROS ultimately damage host DNA, proteins and membrane (Weis 2008; Roth 2014). Moreover, extreme light intensity can be directly damaging for corals and cause a cascade of events that occur from photo-oxidative damage (Roth 2014). Under ambient conditions (i.e. not heat and light stressed), Symbiodiniaceae absorbs light that can be (1) used to drive photochemistry, (2) re-emitted as fluorescence, (3) dissipated as heat, or (4) decayed via the chlorophyll triplet state (Weis 2008; Brodersen et al. 2014; Roth 2014; Roth et al. 2015). Experiments have shown that Symbiodiniaceae in scleractinians, under typical irradiances at shallow coral reefs (640 μmol photons m^−2^ s^−1^), dissipate 96% of the energy and use only 4% of absorbed light energy for photosynthesis (Brodersen et al. 2014). Therefore, in the coral-algae symbiotic association, Symbiodiniaceae harvest sunlight for photosynthesis and dissipate excess energy that can prevent -light induced -oxidative stress (Brodersen et al. 2014; Roth 2014; Roth et al. 2015).

The earliest suggestion of dinoflagellates being present in antipatharian tissues comes from the *Report on the Antipatharia collected by H*.*M*.*S. Challenger* by Brook (1889). A few years later, in *The Antipatharia of the Siboga Expedition* report, van Pesch (1914) documented six species containing what he referred to as ‘symbiotic Algae’ ranging from 7-10 µm in diameter in the gastrodermis. He only reported observing these cells in six out of the thirty species examined in this report (van Pesch 1914). With no more empirical studies or reports for many decades and due to antipatharians ability to thrive at abyssal depths and low-light environments, they were assumed to lack Symbiodiniaceae – commonly referred to as ‘being azooxanthellate’ (Grigg 1965; Wagner et al. 2011a). Moreover, a few later reports using molecular techniques did not find dinoflagellates in the antipatharian species examined. For instance, Grigg (1964) reported dinoflagellates absence in *Antipathes grandis* VERRILL, 1928, from Hawaii after conducting intense morphological studies on the species. Using dinoflagellate-specific primers and spectrophotometric methods that detect dinoflagellate chlorophyll absorbance patterns, Santiago-Vázquez et al. (2007) determined the absence of the symbiotic microalgae in the species *Stichopathes luetkeni* (BROOK, 1889) (formerly called *Cirrhipathes lutkeni*).

However, in accordance with historical suggestions, two more recent studies have confirmed the presence of Symbiodiniaceae in various antipatharian species. Wagner et al. (2011a) conducted a histological analysis of fourteen antipatharian species collected between 10 and 396 m depth from Hawaii and Johnston Atoll. They reported low densities (0–92 cells mm^-3^) of Symbiodiniaceae cells inside antipatharian gastrodermal tissues, suggesting that the dinoflagellates are endosymbiotic. Additionally, dinoflagellates sequences retrieved from antipatharians samples confirmed the presence of Symbiodiniaceae from the genera *Cladocopium, Gerakladium* and *Durusdinium*. Wagner et al. (2011) concluded that the endosymbiotic dinoflagellates had no significant role in the ‘nutrition’ of the different species examined and suggested more research to determine whether the association could be parasitic. The authors’ conclusion was based on the low density of microalgae cells within the antipatharians and their presence in colonies at depths where light penetration does not enable photosynthesis. They did not find a pattern in the recovery of Symbiodiniaceae types from colonies of the same antipatharian species; therefore, the authors suggested that endosymbiont acquisition occurred opportunistically and was not host-specific. The second recent study documenting Symbiodiniaceae in antipatharians was conducted on a single species of the genus *Cirrhipathes* from Indonesia. Bo et al. (2011) sampled two colonies at 38 m and one at 15 m, and found evidence of abundant (∼10^7^ cells cm^−2^) Symbiodiniaceae cells in the corals gastrodermis. They identified two genera, *Cladocopium* and *Gerakladium*, the latest commonly found in association with clinoid sponges. The study concluded that a mutualistic endosymbiosis existed based on the presence of the dinoflagellates inside the antipatharian gastrodermis, a symbiosome surrounding the microalgal cell and evidence of its division inside the host.

Symbiotic associations with Symbiodiniaceae are evolutionarily conserved in all six orders in the subclass Hexacorallia. Antipatharia was the only order in this subclass that was considered the exception before Bo et al. (2011) and Wagner et al. (2011a) presented evidence that it is not. However, these two studies, principally based on the presence and abundance of the Symbiodiniaceae cells and their location in the host tissue led to different conclusions. In both studies, the number of species-specific samples examined was limited. The present study took place within the current framework of limited knowledge, with the objective to bring new insights about Symbiodiniaceae occurrence and density in antipatharians, and to expand the geographic range of studied species. We investigated the presence, abundance, location and identity of Symbiodiniaceae in two antipatharian species -*Cupressopathes abies* (LINNAEUS, 1758) and *Stichopathes maldivensis* COOPER, 1903, representing two different morphologies from shallow and mesophotic reefs of SW Madagascar. A systematic sampling protocol was followed in order to elucidate potential patterns related to coral species, depth and location in the colony. An integrative methodological approach combining microscopy and molecular techniques was developed. Altogether, this study aims to contribute to our understanding of the ecology of the highly important antipatharian corals.

## 2. Materials and Methods

### 2.1. Site description

Sample collection took place near the Great Reef of Toliara (GRT), in SW Madagascar. The GRT is a barrier reef almost 20 km long and 2 km wide, surrounded by two freshwater rivers at its northern (Fiherenana River) and southern (Onilahy River) extremities (Figure 1a). Relevant coral communities -including scleractinians and antipatharians – were first documented in the area on shallow (>30 m depth) and mesophotic reefs (at depths between 35 m and 55 m) in the 1970s (Pichon 1978). More recent studies documented severe reef degradation due to fisheries, pollution, and heavy sedimentation derived from nearby river deposition (Harris et al. 2010; Todinanahary et al. 2016, 2018), as well as coral bleaching episodes (Gudka et al. 2018). Antipatharian samples were collected in November-December 2018 at shallow (20 m depth, 23°20.978’ S, 43°36.885’ E – Site 1) and mesophotic (40 m depth, 23°21.345’ S, 43°36.348’ E – Site 2) reefs (Figure 1a). Figure 1b shows the annual median of the diffuse attenuation coefficient for downwelling irradiance at 490 nm in m^-1^ (K_d_ 490) for the year 2018 in the region obtained from satellite data (MODIS-Aqua). These remote sensing ocean colour coefficient values -in the blue-green spectra range -are used as proxies for water turbidity (Zhang and Fell 2007).

**Figure 1.**
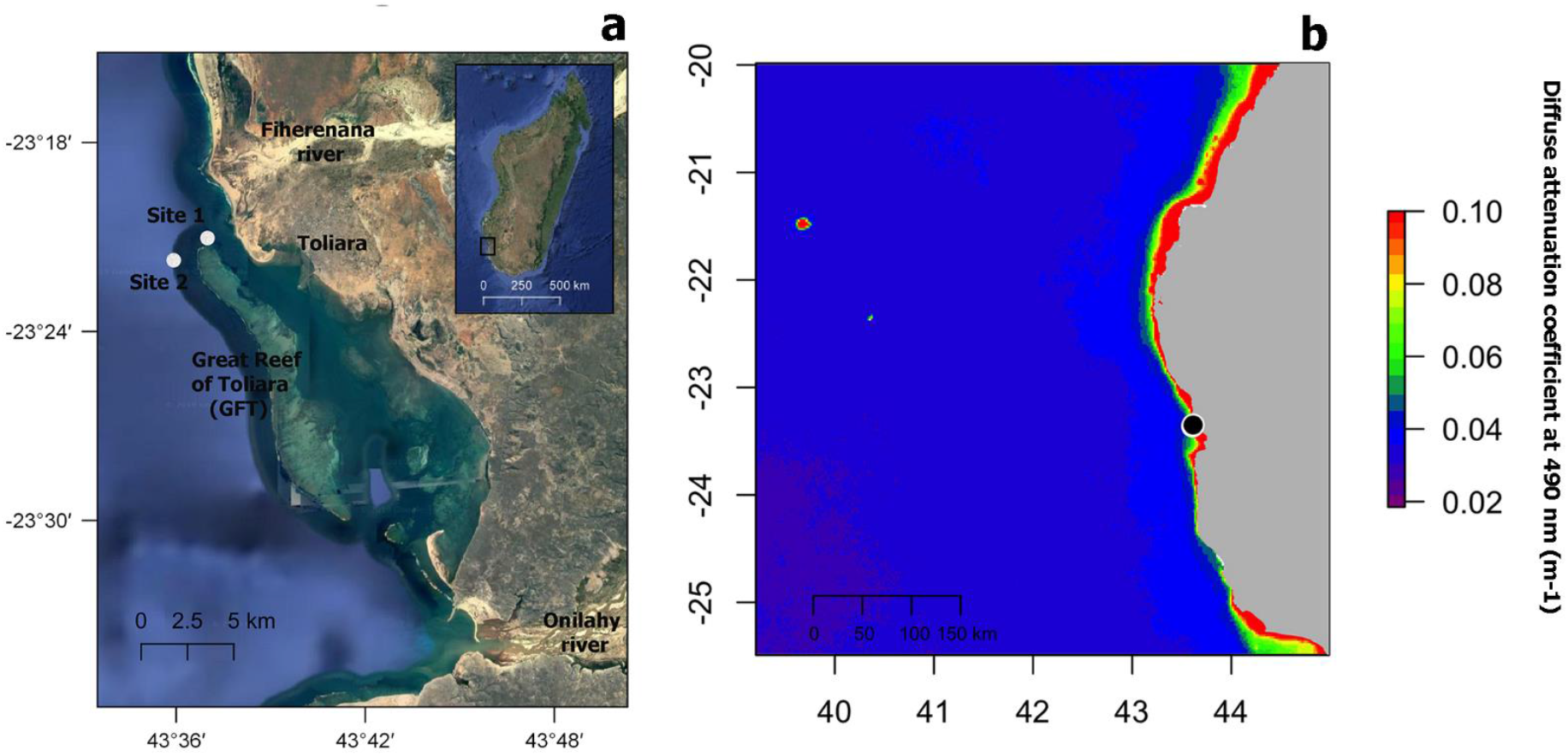
(**a**) Map of study sites 1 and 2 (white circles) near the Great Reef of Toliara (GRT), in SW Madagascar. (**b**) Annual median of the diffuse attenuation coefficient for downwelling irradiance at 490 nm in m-1 (K_d_ 490) for the year 2018 in the region (black circle locates GRT). Satellite data derived from MODIS-Aqua accessed through OceanColor (https://oceancolor.gsfc.nasa.gov). Increasing coefficient values indicate higher water turbidity.

### 2.2. Samples collection

Two antipatharian species, which represent two of the most abundant species at sites 1 and 2, the bottle-brush-like *Cupressopathes abies* (Figure 2a) and the whip-like *Stichopathes maldivensis* (Figure 2b), were sampled. Six colonies of each species were sampled in November-December 2018 at 20 m depth and five colonies of each at 40 m depth (permit no. 089/19/MESipReS). Colony size ranged between 25-35 cm in height for *C. abies* and 200-250 cm in height for *S. maldivensis*. It has been shown that Symbiodiniaceae abundance and clade can vary at the intra-colony level (Meistertzheim et al. 2019) – therefore, three samples of 3-4 cm long were taken for each colony at the top, the middle and the base, making a total of 66 samples from 22 separate colonies. Half of every sample was preserved in 100% ethanol for molecular analysis. For all morphological analyses, the other half was fixed in 3% glutaraldehyde buffered with 0.1M sodium cacodylate before rinsing and storing them in 70% ethanol. Due to the minute size of their polyps (∼0.7 mm, Figure 2a), samples of *C. abies* were placed in a 5 g/L magnesium chloride solution buffered in filtered sea water prior to fixation to relax the polyps and the tentacles, to facilitate further morphological analysis.

**Figure 2.**
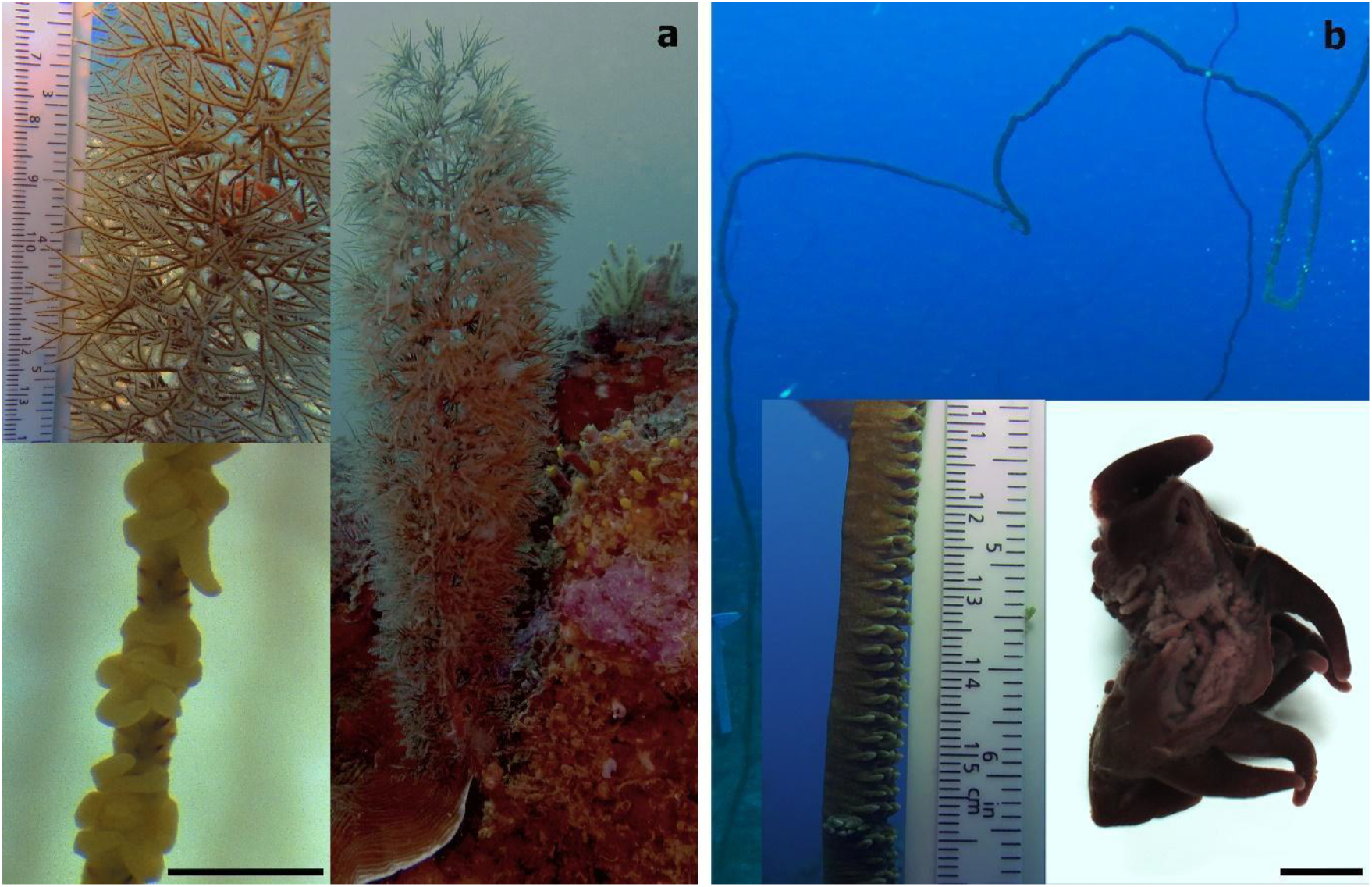
(**a**) Bottle-brush-like *Cupressopathes abies* (LINNAEUS, 1758), and (**b**) the whip-like *Stichopathes maldivensis* COOPER, 1903, with bottom right image showing a section of almost two polyps. Scale bars of polyp images = 1 mm.

### 2.3. Isolation of Symbiodiniaceae cells

From all 66 samples preserved in 100% ethanol, 1 cm fragments were cut. They included about two polyps of *S. maldivensis* and about eight polyps of *C. abies*, an estimated 20 mg of tissue for both species. Following methodology adapted from Zamoum and Furla (2012) to isolate Symbiodiniaceae cells from cnidarians, samples were placed in 1.5 ml tubes with 500 μl of a 2M sodium hydroxide solution. These were incubated at 37 °C for 1 h and vortexed at a medium speed (using an Analitik Jena Tmix homogenizer) at the same temperature for about 2 h until complete lysis of the antipatharian tissues. Tubes were centrifuged at 8,000 rpm for 3 min and the supernatant was discarded, then 8 μl of the pellet was resuspended in 7 μl of ultra-pure (Milli-Q) water and vortexed. One drop of lugol was added to the solution to re-stain the cells that might have lost pigments after preservation. Counts were done using a glass haemocytometer and an Axioscope A1 (Zeiss) light microscope. Microalgae cells in all nine squares (1 mm^2^ each) of the counting chambers were considered to estimate the density of cells per μl, and four chambers were counted per sample. Isolation protocol was also carried out on fresh and preserved (100% ethanol) samples of the scleractinian coral *Seriatopora hystrix* to corroborate efficiency (Figure 3). To test for differences in microalgae density between antipatharian species, depths, and colony region (top, middle or bottom part), a quasipoisson generalised linear model (GLM) was fitted using the ‘stats’ package. The power to detect differences between depths was calculated using the function ‘power.t.test’ in the ‘stats’ package. Analyses were conducted in R (R Team 2019).

**Figure 3.**
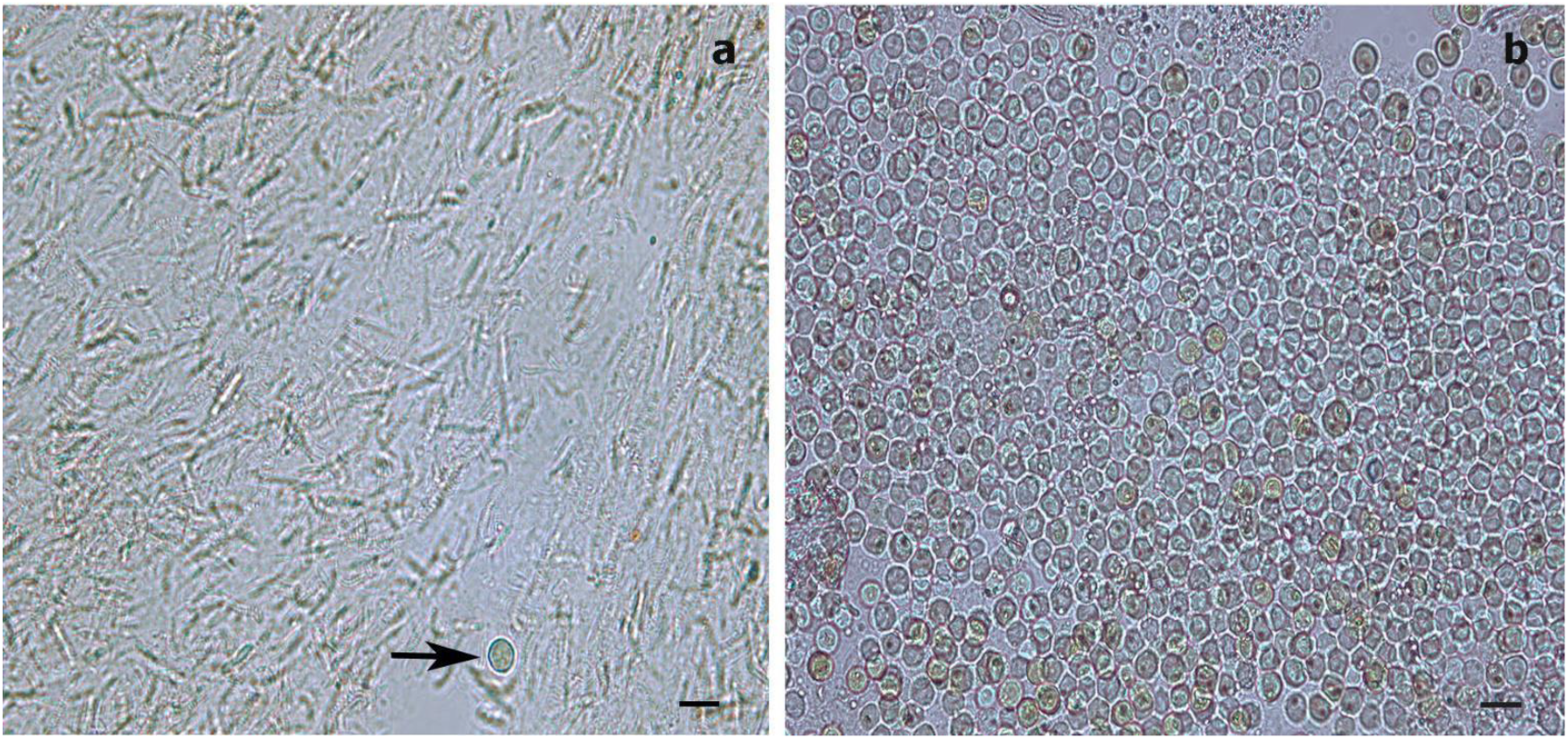
Light microscopy images of Symbiodiniaceae-like extracts obtained through the sodium hydroxide isolation method. (**a)** Extract from the antipatharian *Cupressopathes abies* showing only one Symbiodiniaceae-like cell (arrow) and abundant cnidocytes. (**b**) Extract from the scleractinian *Seriatopora hystrix* depicting numerous Symbiodiniaceae-like cells, corroborating the efficiency of the isolation method. Scale bars = 10 μm.

### 2.4. Microscopy

#### a) Histology

Histological sections were made to determine the presence and location of Symbiodiniaceae cells within the corals tissue. About 1.5 cm of each sample was dehydrated using increasing concentrations of ethanol before being soaked in butanol at 60 °C for 24 h. These were then embedded in liquid paraffin. Serial sections of 7 μm were cut using a Microm HM 340E (Zeiss) microtome and stained with a Masson’s trichrome. The sections were observed and photographed using an Axioscope A1 (Zeiss) light microscope and Axiocam 305 camera.

#### b) Transmission electron microscope (TEM)

Sub-sample sections were made from the top, middle and bottom part of one colony from 20 m depth and one from 40 m depth of each species. For *S. maldivensis*, sections of the tentacles and of the oral cone were analysed separately due to the large size of the polyps (∼6 mm, Figure 2b), making a total of 18 sub-samples examined. These sections of about 1 cm were post-fixed for 1 h at room temperature with 1% osmium tetroxide, in a 0.1 M sodium cacodylate and 2.3% sodium chloride buffer. They were rinsed several times in the same buffer and dehydrated in increasing concentration series of ethanol. Samples were then placed in Spurr resin overnight before polymerisation at 70 °C for 24 h. Ultrathin sections between 50-70 nm were cut on a Leica Ultracut (UCT) microtome equipped with a diamond knife and collected on formvar-coated copper grids. These were stained with uranyl acetate and lead citrate, and observed with a LEO 906E (Zeiss) transmission electron microscope.

#### c) Scanning electron microscope (SEM)

Middle sections of one colony from 20 m depth and one for 40 m depth of each species were analysed in SEM. The four samples of about 1.5 cm, stored in 70% ethanol, were dehydrated as for TEM analysis. Sub-samples were also made in order to be frozen with liquid nitrogen before being cut randomly. The eight sub-samples were dried in a critical-point dryer using CO_2_ as the transition fluid (Agar Scientific Ltd.) before being mounted on aluminium stubs and coated with gold in a JFC-1100E (JEOL) sputter coater. These samples were observed and photographed with a JSM-7200F (JEOL) scanning electron microscope.

### 2.5. Molecular analyses

Middle parts of the colonies from both species, and from both depths (preserved in 100% ethanol) were cut into 8 mm fragments. Total genomic DNA of each sample was extracted using QIAGEN DNeasy Blood & Tissue Kit using the manufacturer’s protocol. Concentration and quality of DNA were examined at 260 nm using a spectrophotometer (DeNovix). The internal transcribed spacer-2 (ITS2) region of the dinoflagellate ribosomal DNA (rDNA) was amplified using the primers ITS-DINO (5′ GTGAATTGCAGAACTCCGTG 3′) and ITS2-REV2 (5′ CCTCCGCTTACTTATATGCTT 3′) following conditions described by Stat et al. (2009). A second set of primers, SYM_VAR_5.8S2 (5’ GAATTGCAGAACTCCGTGAACC 3’) and SYM_VAR_REV (5’ CGGGTTCWCTTGTYTGACTTCATGC 3’) were used for PCR amplification (ITS2 region) following conditions described by Hume et al. (2018). This second set of primers was also used because it was found to perform superior to other ITS2 primer sets tested on a range of Symbiodiniaceae ITS2 rDNA (Hume et al. 2018). The presence of amplicons was checked in a 2% agarose gel in Tris-Borate-EDTA buffer. A scleractinian coral DNA extract of the genus *Acropora* from Toliara (Madagascar) was used as positive control.

## 3. Results

### 3.1. Dinoflagellate cells count

Isolation of Symbiodiniaceae-like cells with sodium hydroxide was the procedure used to analyse all 66 samples. This protocol enables efficient isolation of the microalgal cells if present within the coral tissues, and it is also an efficient way to isolate cnidocytes (Figure 3). To ease comparison with one of the two previous studies (Wagner et al. 2011) we will report results in the equivalent cells mm^-3^ instead of cells μl^-1^. For both antipatharian species and depths, dinoflagellates cells density ranged between 0-4 cells mm^-3^. No significant difference was detected based on region on the colony using the quasipoisson GLM, therefore the region factor was removed from the model. The average dinoflagellate cell density from the three regions of each colony was calculated to get a mean cell density per colony for further analyses. Microalgae density was different between the two species (*t=*2.36, *p=*0.029; Table 1), with a mean density of 0.037 ± 0.013 cells mm^-3^ (mean ± SE) for *C. abies* and 0.121 ± 0.034 cells mm^-3^ for *S. maldivensis*. No significant difference was detected in Symbiodiniaceae-like density between depths (*t=-*1.577, *p=*0.131; Table 1). The statistical power to detect a significant difference in cell density between depths for each species was evaluated. Results show a low power of 0.26 (Type II error rate 74 %) for *S. maldivensis*, and 0.06 (Type II error rate 94 %) for *C. abies* to detect differences between depths.

**Table 1.** Results of quasipoisson GLM test for differences between species and depths. The intercept represents *C. abies* dinoflagellate cell density. Significant *p-value* (*p <0*.*05*) are shown in bold.

| Factor | Estimate | Standard Error | t-value | p-value |
| --- | --- | --- | --- | --- |
| Intercept | -2.27 | 0.74 | -3.08 | **0.006** |
| *S. maldivensis* | 1.16 | 0.49 | 2.36 | **0.029** |
| Depth | -0.04 | 0.02 | -1.58 | 0.131 |

### 3.2. Morphological analyses

Histological sections were only successfully made for all the S. maldivensis samples. Symbiodiniaceae-like cells in the polyps and gastrovascular gastrodermis of *S. maldivensis* were observed on the sections examined (Figure 4). From most paraffin blocks of *C. abies*, sections of the tentacles and oral cone were not obtained because there is only a thin layer of tissue surrounding the skeleton, and cells likely to be Symbiodiniaceae were not observed on the few successful sections made for *C. abies*. Quantification of these Symbiodiniaceae-like cells was not done from histological sections since they were not obtained for both species. A potential solution for histological analysis of branching antipatharians for future studies, could be by softening the skeletons with a lytic polysaccharide monooxygenases (LPMOs) treatment (Mutahir et al. 2018). Ultrastructural (TEM) observations were successful for all 18 samples examined from both species, and no Symbiodiniaceae-like cells were identified on any of them. Mucous cells, zymogen granules, and the high number of cnidocytes, including spirocysts and b-mastigophores were evident here (Figure 5). Similarly, on SEM examinations round cells of different sizes (3-10 μm) were observed, where the round larger ones (∼8 μm; Figure 6a) could potentially be dinoflagellates cells. Smaller round cells (3-4 μm) were more abundant, which are likely to be the vesicular mucous cells also observed on the ultrastructural analysis (Figure 6b).

**Figure 4.**
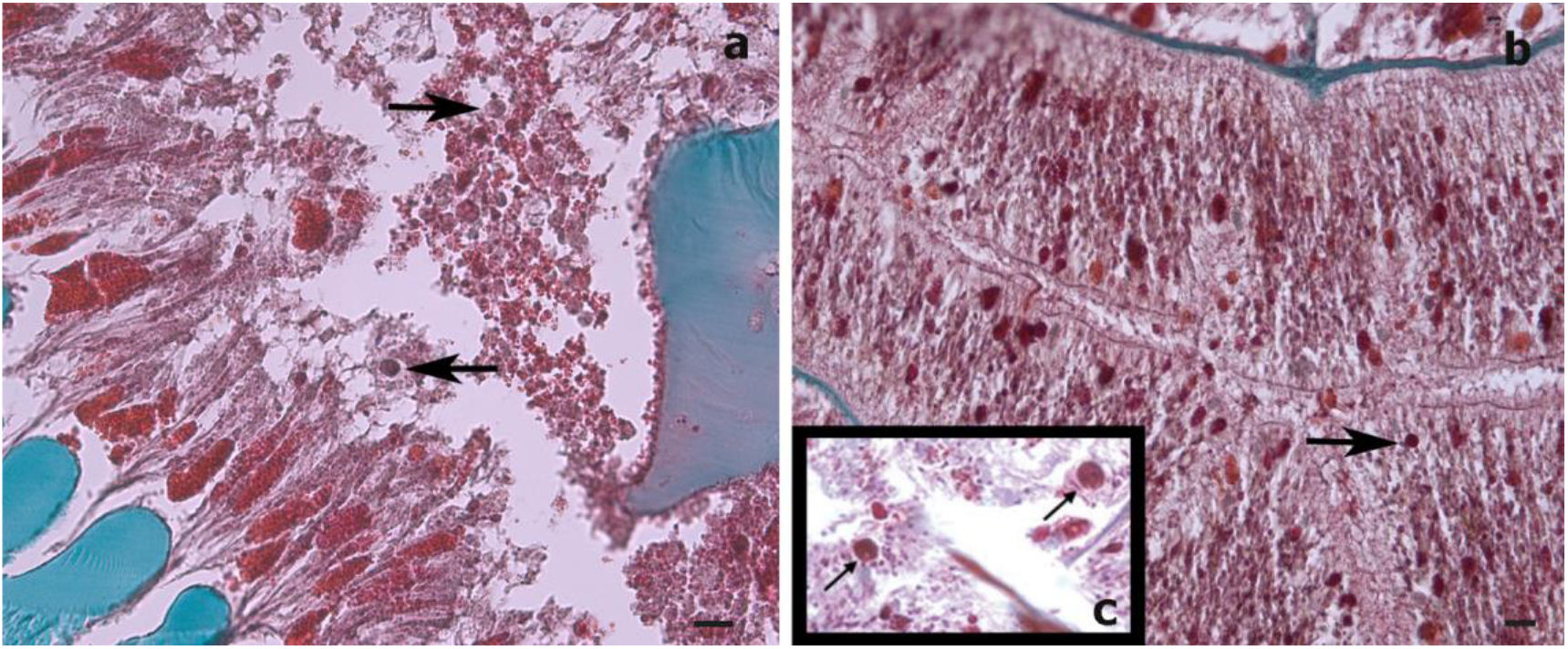
Histological cross-sections of a *Stichopathes maldivensis* polyp. Symbiodiniaceae-like cells (arrows) are observed in the gastrodermis of the gastrovascular cavity (**a**), and in the gastrodermis of a lateral tentacle (**b**). (**c**) Inset from Wagner et al. (2011, Figure 2c) where Symbiodiniaceae cells were identified through histological cross-sections and molecular analysis. Scale bars = 20 μm.

**Figure 5.**
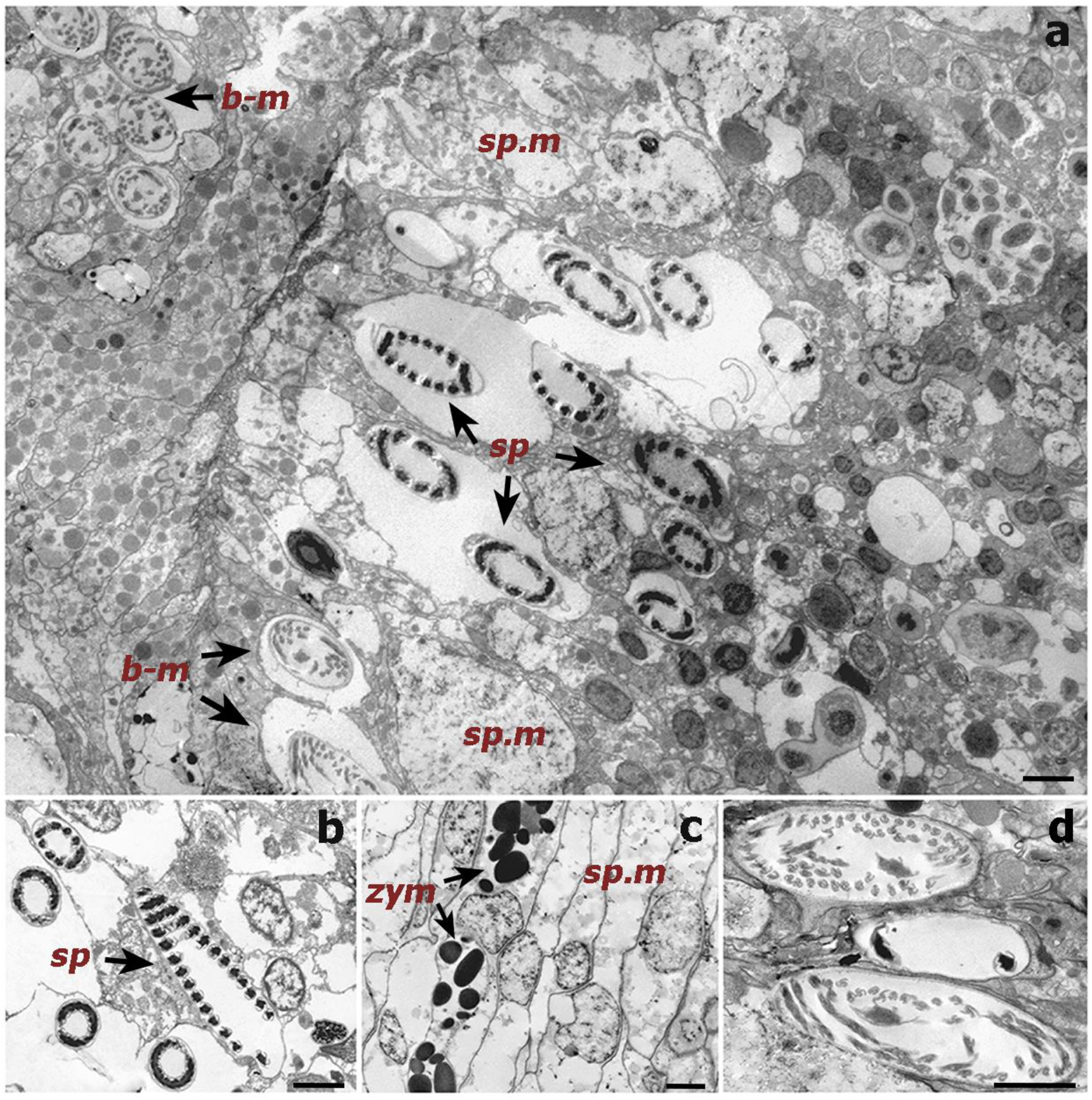
Transmission electron microscope (TEM) images of a cross-section of a *Cupressopathes abies* polyp tentacle. (**a**) Ultrastructural view of numerous cnidocytes: spirocysts [sp] and b-mastigophores [b-m], as well as spumous mucous cells [sp.m]. (**b**) Longitudinal section of mature spirocysts [sp]. (**c**) Close view of zymogen granules [zym] and spumous mucous cells [sp.m] inside the gastrodermis. (**d**) Cross-section of mature b-mastigophores. Scale bars = 2 μm.

**Figure 6.**
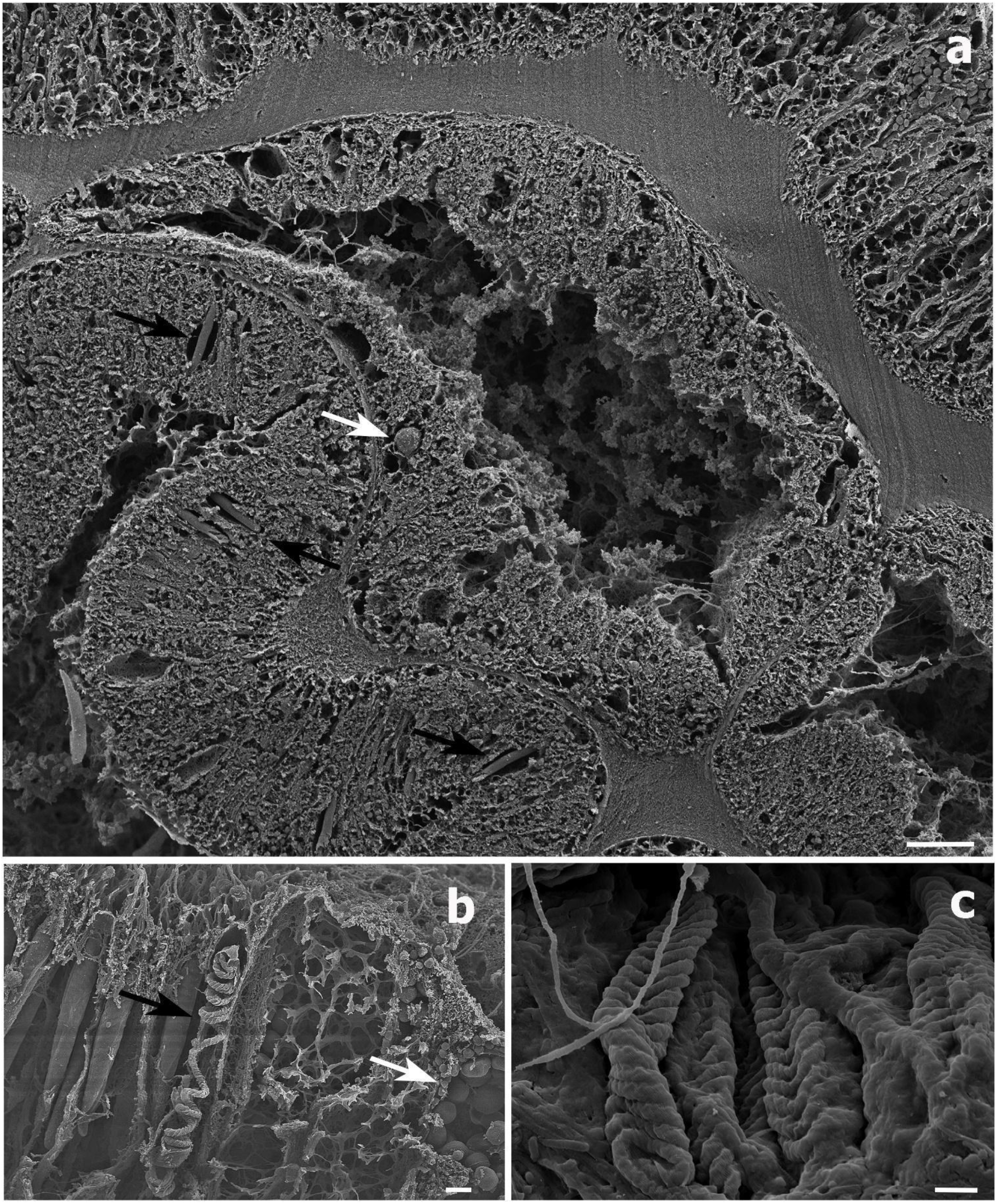
Scanning electron microscope (SEM) images of a *Stichopathes maldivensis polyp*. (**a**) A round -potentially dinoflagellate -cell of about 8 μm inside the gastrodermis of the polyp gastrovascular cavity (white arrow). Numerous b-mastigophores can be observed in the ectoderm (black arrows). (**b**) Smaller round cells (3-4 μm) that are likely to be mucous cells (white arrow) in the polyp ectoderm. Image also shows several b-mastigophores aligned -one has a broken capsule which exposes the cnidocyte tubule (black arrow). (**c**) Closer view of spirocysts in a polyp tentacle ectoderm. Scale bars: a = 10 μm, b = 2 μm, c =1 μm.

### 3.3. Symbiodiniaceae molecular analysis

No ITS2 sequence of Symbiodiniaceae succeeded to be amplified from any of the antipatharian DNA extractions. Only the positive control (scleractinian coral DNA extraction) produced a band on the agarose gel. This was the case for both sets of primers used.

## 4. Discussion

### 4.1. Overall analysis results

Climate change is causing widespread disruptive effects on coral reefs via heat and light induced oxidative stress, which is altering the endosymbiotic relationship between Symbiodiniaceae and scleractinian corals. These concerns were latent with respect to the hitherto assumed ‘azooxanthellate’ antipatharian corals. This changed with recent findings of high densities (∼10^7^ cells cm^−2^) of microalgae cells within the gastrodermis of three shallow antipatharian whip-like colonies (*Cirrhipathes* sp.) (Bo et al. 2011). A methodological study with a higher number of samples was appropriate to assess microalgae densities in a greater number of antipatharian colonies, and to identify possible patterns regarding species, colony morphology and depth. This study shows a very low density (0-4 cells mm^-3^) of Symbiodiniaceae-like cells in both, the whip-like *S. maldivensis* and the bushy *C. abies* at shallow and mesophotic reefs. These findings align with other historical observations (Brook 1889; van Pesch 1914) and more recent studies (Grigg 1964; Santiago-Vázquez et al. 2007; Wagner et al. 2011a). The Symbiodiniaceae-like cells observed from SEM and histological examinations from thin sections of *S. maldivensis*, were located inside the coral gastrodermis (Figures 5, 6). This endosymbiotic association was also suggested by Wagner et al. (2011a) from histological sections analysis, and was corroborated by Bo et al. (2011) through ultrastructural analysis.

No Symbiodiniaceae ITS2 sequences were amplified in this study, most likely due to the extremely low density of dinoflagellates. A potential solution for future studies, if dealing with low abundance of Symbiodiniaceae cells, could be Fluorescence In Situ Hybridization method (FISH) (Figueroa et al. 2018) or qPCR (Saad et al. 2020). From the other two studies where identification was possible, different genera (*Cladocopium, Gerakladium* and *Durusdinium*) were reported to be in association with the antipatharians (Wagner et al. 2011; Bo et al. 2011). This plasticity, even at the intra-colony level, was evidenced in more recent studies on scleractinian corals (Meistertzheim et al. 2019) and is believed to be environmentally driven. Moreover, the same Symbiodiniaceae species may be mutualistic in one host context but opportunistic in another (Pettaya et al. 2015). Therefore, the identity of the dinoflagellates is not sufficient to determine the type of symbiosis. Further studies will be necessary to determine the type of association between Symbiodiniaceae and antipatharians, particularly when microalgae abundance is relevant such as in Bo et al.’s (2011) findings.

Despite not being able to corroborate the dinoflagellates identity by means of molecular analysis, the Symbiodiniaceae-like cells observed on histological sections resemble to a great extent the cells from the other two studies in which their identity was confirmed (Bo et al. 2011; Wagner et al. 2011, Figure 4). In addition, we applied the sodium hydroxide isolation protocol to a scleractinian coral fragment, from where it was clear that these cells are Symbiodiniaceae (Figure 3). Nevertheless, because of inconclusive molecular results we will refer to the cells as Symbiodiniaceae-like.

### 4.2. Dinoflagellate density difference between species

Density estimates showed that colonies of the whip-like *S. maldivensis* had significantly more Symbiodiniaceae-like cells within their tissues compared to *C. abies* regardless of depth (0.084 mean cells mm^-3^ density difference, Table 1). However, due to the extremely low densities of dinoflagellates in both species in general, it is difficult to determine the biological relevance of such difference. From the prior two studies that estimated Symbiodiniaceae densities in antipatharians, only Wagner et al. (2011) examined multiple species and different colony morphologies. The authors present results of dinoflagellates cell density estimates from five different species, two whip-like and three branching from Hawaii. The branching species, *Antipathes griggi*, had the highest density (0-92 cells mm^-3^), although microalgae cells were observed only in one out of the eight colonies examined from this species. Therefore, no obvious patterns regarding species and morphologies can be determined.. Variability was also observed in this study on antipatharians from Toliara, Madagascar -which is the first study that has examined several colonies of the same species at two different depths. From the eleven colonies examined of each species, only four colonies of *C. abies* and seven of *S. maldivensis* contained microalgae. In addition, no difference was found between the three different regions (top, middle and base) within each colony. This suggests that the high intra-specific variability previously recorded by Wagner et al. (2011a) is unlikely to be caused by variation of the sample region within the Hawaiian colonies. The second study presenting microalgae cell densities in antipatharians -from Bunaken, Indonesia (Bo et al. 2011) -only analysed three samples of the whip-like *Cirrhipathes* sp. and did not specify the region within the colony sampled. From this limited existing information, it seems plausible that environmental conditions might be able to explain the higher cell density patterns, rather than the antipatharian species or their morphology.

### 4.3. Dinoflagellate density difference between depths

No significant difference in dinoflagellate density was found between shallow and mesophotic depths. Despite the low statistical power and non-significant results, the mean dinoflagellates density at 20 m depth was higher than at 40 m depth for both species -with a difference between depths of 0.200 and 0.016 mean dinoflagellates cells mm^-3^ for *S. maldivensis* and *C. abies*, respectively. However, interpretation of such small differences is not possible. Differences in microalgae density can be inferred from previous studies. For instance, the sample of *A. griggi* from Hawaii -containing the higher density in that study -was collected from shallow reefs (24 m depth) (Wagner et al. 2011), and of all the 14 samples from different colonies were cell density was reported, the highest densities were from the two colonies sampled in shallow reefs (Wagner et al. 2011). The samples of *Cirrhipathes* collected in Bunaken, Indonesia (Bo et al. 2011) were from two different depths: 15 m (reef wall) and 38 m (reef slope). It was not specified if there was any difference in Symbiodiniaceae density between the samples from these two depths. However, K_d_ 490 values at 15 m and 38 m depth in Bunaken have been calculated to be very similar (Holden 2002), which indicates similar high water clarity at both depths. This is not the case in Toliara, Madagascar, where K_d_ 490 values show high turbidity at 20 m depth (Figure 1b), and due to sedimentation derived from river runoff, light penetration at 40 m depth is likely to be considerably reduced (which was noticeable during field work). A higher number of free living dinoflagellates is expected at shallow water environments -where they can photosynthesise -even though Symbiodiniaceae’s depth range extends down to 396 m (Wagner et al. 2011). Nonetheless, this does not mean that higher densities of Symbiodiniaceae will inevitably be in association with shallow antipatharians. This is because the uptake of dinoflagellates in the coral-algal symbiotic association is believed to be controlled by the coral host (Davy et al. 2012; Barott et al. 2015).

### 4.4. Potential environmental role on the Symbiodiniaceae-antipatharians association

Understanding the benefits antipatharians could gain in a mutualistic symbiotic association with the Symbiodiniaceae requires further physiological studies (e.g. metabolomics and photobiology). The process of endosymbiotic microalgae supporting corals metabolism is better known from scleractinians. Even if not fully understood, it is known that photosynthetically fixed carbon is translocated to the host supporting its growth and respiration (Davy et al. 2012). It also has been postulated that symbiotic dinoflagellates may stimulate scleractinians calcification by either absorbing CO_2_ or releasing O_2_, or they might produce organic molecules that buffer the protons during precipitation of the scleractinian calcium carbonate skeleton (Davy et al. 2012; Barott et al. 2015). Antipatharians have a proteinaceous skeleton not a calcareous one. Also, they are known to be effective phytoplankton, zooplankton and dissolved organic matter (DOM) feeders (Terrana et al. 2019; Rakka et al. 2020), armed with abundant and diverse cnidocytes (Goldberg and Taylor 1989; Figures 3,5, 6). It cannot be ruled out that the high abundance of Symbiodiniaceae in antipatharians found in other studies is associated with limited food supply. However, antipatharians have the ability to exploit high quantities of shortly available pray (Rakka et al. 2020). This explains the higher variety and abundance of cnidocytes in antipatharians compared to scleractinians (Figures 3, 5, 6), and the common findings of low or none Symbiodiniaceae within antipatharians. Therefore, if a mutualistic symbiotic association is possible (Bo et al. 2011), the microalgae might be supporting antipatharians with other processes. These processes could be directly related to an important environmental factor at shallow depths: high light irradiance, where higher densities of dinoflagellates in antipatharians have been reported by Bo et al. (2011) and Wagner et al. (2011).

Light absorbed by the microalgae is known to be used to drive photochemistry, then be re-emitted as fluorescence, dissipated as heat, or decayed to produce ROS (Weis 2008; Roth 2014). On a sunny day, Symbiodiniaceae in shallow corals dissipate four times more light energy than is used in photosynthesis (Brodersen et al. 2014; Roth 2014). Consequently, in the coral-algae symbiotic association on ambient conditions, Symbiodiniaceae harvest sunlight for photosynthesis and dissipate energy excess to prevent oxidative stress (Weis 2008; Roth 2014). Antipatharians are most commonly found in low-light environments (Grigg 1964, 1965; Grange and Singleton 1988; Wagner et al. 2011a, 2012). Therefore, the high densities of microalgae from antipatharian colonies at shallow reefs that have been reported – particularly the *Cirrhipathes* colonies from Bunaken, Indonesia and *A. griggi* from Hawaii – could be related to ultraviolet radiation. Accordingly, it might be suitable to investigate whether antipatharians exposed to higher light radiations (compared with environments in which they usually settle) enhance their ability to develop the symbiotic relationship with microalgae to get assistance with light absorption and utilization, and dissipate excess energy to prevent oxidative stress and/or photo-oxidative damage. To draw any conclusions in this regard and to investigate any potential climate change hazards, more physiological studies on antipatharians -spanning different regions, species and depths -are needed. Yet, it might be appropriate to prioritize those in colonies exposed to high levels of light radiation, where the highest microalgae densities have been documented.

## 5. Conclusions

This study represents the first methodological and integrative approach to investigate the presence and estimate the density of Symbiodiniaceae in antipatharians. Symbiodiniaceae-like cells were present within antipatharian tissues of *S. maldivensis* and *C. abies* from SW Madagascar. However, the overall density of the macroalgae cells in the two antipatharians species from both shallow and mesophotic reefs was very low. This low density aligns with past observations and studies, and suggests that high Symbiodiniaceae densities are not characteristic in antipatharians. These findings are relevant in the context of ‘bleaching’ events threatening shallow and mesophotic reefs, and suggest that antipatharians might be less prone to such disturbances. Nevertheless a trend of higher dinoflagellate densities in antipatharians from shallow reefs was noticed in the present study, which aligns with previous studies where high densities of Symbiodiniaceae have been reported. Further physiological studies, particularly on colonies exposed to high ultraviolet radiations are necessary to better understand these trends. In addition, more studies on antipatharians and their association with dinoflagellates could increase our understanding of the mechanisms involved in the coral-algae endosymbiotic association.

## Author contributions

Conceptualization: Erika Gress and Igor Eeckhaut. Writing – original draft and figures preparation: Erika Gress. Data curation: Erika Gress, Mathilde Godefroid, and Jonathan Richir. Formal analysis: Erika Gress. Funding acquisition: Igor Eeckhaut, Philippe Dubois and Lucas Terrana. Methodology: Erika Gress, Igor Eeckhaut and Lucas Terrana. Writing – review & editing: Igor Eeckhaut, Mathilde Godefroid, Philippe Dubois, Jonathan Richir and Lucas Terrana.

## Acknowledgements

We thank: All the members at the Marine Organisms Biology and Biomimetics (BOMB) Laboratory, in Belgium, for their assistance during samples analyses. Nicolas Sturaro, University of Liege, for help with samples collection. Members at the University of Toliara, Madagascar for support during field work. Liz Tynan, James Cook University, for editorial feedback. Figure 1b created on R (R Team 2019) with an adapted script provided by J.C. Fischer, University of Bayreuth and Dr. A. Wiefels, University of Reunion Island. We are very grateful to Dennis Opresko, Smithsonian Institute, for continuous support and for sharing his vast knowledge on antipatharians.

